# Single generation exposure to a captive diet: a primer for domestication selection in a salmonid fish?

**DOI:** 10.1101/2020.01.24.919175

**Authors:** Shahinur S. Islam, Matthew C. Yates, Dylan J. Fraser

**Author notes:** Both authors contributed equally (Shahinur S. Islam, Matthew C. Yates). Author to whom correspondence should be addressed: Dylan J. Fraser, Department of Biology, Concordia University, Montreal, QC, Canada.

## Abstract

Millions of wild animals in captivity are reared on diets that differ in their uptake and composition from natural conditions. Few studies have investigated whether such novel diets elicit unintentional domestication selection in captive rearing and supplementation programs. In highly fecund salmonid fishes, natural and captive mortality is highest in the first few months of exogenous feeding. This high early mortality might be a potent driver of unintentional selection because wild fish normally forage on live prey whereas they are fed almost exclusively pellet feed in captivity: fish that do not adapt pellet feed well under captive conditions experience reduced growth and/or die. We tested this hypothesis by generating a large number of families from F_1_ captive and wild fish originating from the same three populations and then rearing them each on pellet and natural, live, drifting feed for three months at the beginning of exogenous feeding. We found that captive fish of every population grew faster than wild fish in all diet treatments. Populations exhibited an idiosyncratic response to diet treatment, with two populations exhibiting faster growth on a pellet diet versus the natural diet but another population exhibiting similar growth in both diet treatments. Fish exposed to a natural diet also exhibited higher survival relative to those given a pellet diet. Captive and wild fish did not differ in survival, regardless of population of origin. Overall, we found evidence that rapid domestication selection associated with a single generation exposure to a novel captive diet generates genetically-based changes to individual fitness (e.g., growth and survival) in a wild fish.

## Introduction

Captive rearing is routinely implemented in biodiversity conservation to prevent species extinctions. Yet captive and natural environments invariably differ in selective pressures, which cause genetic changes to captive phenotypes that reduce fitness in nature and that can ultimately hinder wild population persistence and recovery (Frankham, 2008; Fraser, 2008; Bowlby & Gibson, 2011). Such maladaptive phenotypic and genetic changes have been observed in as few as one or two captive generations (Araki et al., 2007; Christie et al., 2012, 2016; Fraser et al., 2019) and can also have carry-over fitness effects in nature (Araki et al., 2009; Evans et al., 2014; Clarke et al., 2016; Edelaar & Bolnick, 2019).

The diet and food uptake of captive animals frequently deviates from what is normally experienced in the wild. For example, many captive bird and fish species receive novel commercial pellet feeds (Carciofi et al., 2006; Brightsmith, 2012; Lall and Dumas, 2015; Li & Robinson, 2015), and many carnivorous mammals are fed combinations of animal proteins (Kyriazakis et al., 1991; Mayntz et al., 2009; Erlenbach et al., 2014; Hewson-Hughes et al., 2016). Although these diets try to emulate the nutritional requirements of animals, they are often novel in texture and shape, they may not match such requirements precisely, and they may not stimulate active feeding behaviours that are required by wild animals to seek out and obtain prey (Mayntz et al., 2005; Benhaim et al., 2013; De Mestral & Herbinger, 2013). As a result, certain captive individuals may more readily adopt commercial diets than others (Le Boucher et al., 2012; Allan et al., 2018), with consequences for subsequent growth, body condition, reproductive output, and survival. However, little empirical research has investigated how novel captive diets might elicit unintentional domestication selection in captive reared animals.

Owing to their cultural and economic significance and to the tremendous amount of unique population diversity they harbour, salmonid fishes are some of the world’s most extensively captive-reared vertebrates. Tens of billions of juveniles reared in hatcheries are stocked annually to supplement declining wild populations (Naish et al., 2008; Gozlan et al., 2010). Unintentional domestication selection pervades many salmonid captive supplementation programs because hatchery and wild environments differ in densities, developmental timing, abiotic/biotic conditions and food availability (e.g., Reisenbichler & Rubin, 1999; Fleming & Petersson, 2001; Fraser, 2008; Christie et al., 2016; Hagen et al., 2019). While wild salmonids begin exogenous feeding on natural drifting prey, most captive salmonids are fed on an artificial pellet diet designed to optimize growth, despite simultaneous attempts to minimize unintentional domestication selection in other ways (Ford, 2002; Fraser, 2008; Baskett & Waples, 2012; Clarke et al., 2016).

Few studies have investigated how artificial pellet feed might induce genetic changes in wild salmonid populations that are reared in captivity (Dender et al., 2018; Janisse et al., 2019), even though there is now an extensive literature on how captivity affects salmonid phenotypes, genetic characteristics and individual fitness (Le Cam et al., 2015; Christie et al., 2018; Waters et al., 2018; Fraser et al., 2019). In nature, early juvenile salmonids feed on drifting and benthic invertebrates, and the availability, type and nutrient composition of such prey will depend on habitat characteristics such as a flow rate and substrate (Braithwaite and Salvanes, 2005; Jonsson & Jonsson, 2011; Naslund & Johnsson, 2016). Failure to initiate and maintain effective pellet feeding under higher density captive conditions may result in mortality or reduced growth, and has the potential to lead to subsequent changes to fitness-related traits of wild individuals. Moreover, the diet shifts in captivity can elicit stress in certain individuals (Solberg et al., 2015). Collectively, a novel diet in captivity may elicit widespread, unintentional domestication selection, especially because growth and mortality are often highest in the first few months of exogenous feeding in highly fecund species such as salmonids.

Wild populations might also differ in their responses to captive diets (Woodworth et al., 2002; Reisenbichler, 2004; Fraser et al., 2019). Such differences might be expected initially as populations are rarely coming into captivity from the same genetic/phenotypic ‘starting conditions’ (Fraser et al., 2019). However, this could be followed rapidly by subsequent phenotypic homogenization given the homogeneous nature of captive environments (Morris et al., 2011) and indications of a common ‘cultured phenotype’ arising from domestication selection (Wringe et al., 2016). Furthermore, the rapidity by which phenotypic plasticity might evolve during adaptation to captivity has rarely been investigated, despite the role that plasticity plays in modulating adaptive organismal responses to environmental change (Reed et al., 2011). Plasticity might specifically be selected against in captivity, because such temporally stable environments may not favour its retention (DeWitt & Scheiner, 2004).

Here we tested four hypotheses concerning how a captive diet may generate genetic changes to fitness traits in a salmonid fish (brook trout, *Salvelinus fontinalis*): (i) it can arise after a single generation of captive rearing, (ii) it differs among wild populations; (iii) it homogenizes associated phenotypes; and (iv) it reduces phenotypic plasticity. Trout from three wild populations were captive-reared for one generation on pellet feed only. Once these captive trout matured, we simultaneously generated F_1_ captive families as well as wild families from each of the same three populations by recollecting wild trout gametes. We then compared the early growth and survival of newly feeding captive and wild families in common environmental conditions under two diet regimes: pellet feed and a more “natural” diet of live feed (cultured artemia). If genetic adaptation to captivity was taking place, then F_1_ captive families from each population, whose parents survived and reproduced on novel pellet feed, would be expected to have better growth and survival when fed on a diet of commercial pellets relative to their wild family counterparts with no such ancestral exposure to this diet.

## Materials and Methods

### Study Site and Populations

Brook trout used in this study originated from three wild populations located at Cape Race (CR), Newfoundland, Canada: Upper Ouananiche Beck (OB), Watern Cove River (WN) and Freshwater River (FW) (Fig. S1). Many low-lying streams in CR are inhabited by isolated, pristine and genetically distinct brook trout populations (Wood et al., 2014; Bernos & Fraser, 2016). Despite being separated only 5km apart, the three study populations experience significant spatio-temporal environmental differences (Wood et al., 2014) and differ substantially in adaptive genetic characteristics (Fraser et al., 2014; Wood et al., 2015) and life histories (Hutchings, 1993; Fraser et al., 2019).

### Gamete Collections

In late-October of 2011 and 2014, gametes were collected from wild males and females from OB, WN, and FW. Families generated from 2011 gametes produced the parents of F_1_ captive trout used in this study (details in captive rearing of parental crosses section); 2014 gametes collected from the wild were used to generate our study’s wild families. In both years, electrofishing was used to collect spawning individuals from previously documented spawning sites (dense aggregation of matured broods) (Wood & Fraser, 2015). Mature males and females were placed in a flow-through cage for 24-72 hours until gamete collection took place. Eggs and sperm were collected in 60ml opaque plastic containers and 1.5ml micro-centrifuge tubes, respectively. Gametes were kept in coolers, transported from CR to St. John’s (Newfoundland), flown to Montreal, and crossed at Concordia University within 15 hours from the beginning of gamete collection.

### Captive Rearing of Parental Crosses

Once hatched (spring 2012, see Fraser et al. 2019 for more details), captive trout from the three populations were raised for 20 months under common environmental conditions (temperature, pH, dissolved oxygen, density) and on a diet of aquaculture feed (CoreyAqua) administered *ad libatum* (multiple times daily for six months, once daily afterwards). Temperatures within tanks were allowed to fluctuate seasonally between 3°C and 19°C but were always kept within 0.2-0.3°C of each other (see details in Fraser et al., 2019). In late November 2014, gametes were collected from these captive adults to create F_1_ captive families; in this study, each female was crossed with one male from the same population to generate full-sibling families.

### Experimental Design

Replicates of full-sibling families generated in 2014 (wild or captive origin) were placed randomly in 5.2 cm diameter individual egg containers into two 1000-L recirculation tanks for the entire incubation period up to yolk sac absorption, i.e. the beginning of exogenous feeding. Tubes were covered with a mesh bottom to allow for water circulation; pH was approximately 7.0 and dissolved oxygen was maintained at saturation level. Eggs were kept untouched except for the periodic removal of fungal-carrying eggs during incubation.

After yolk sac absorption, 7-10 families from each of the three populations and two origins (wild or captive), were replicated twice randomly in tubes across two tanks, with four individuals per replicate; one family replicate receiving each diet treatment per tank. Thus, in total, 859 trout (Table S1) were used in the experiment (7-10 families × 3 populations × 2 treatments × 2 sources × 2 replicates × 4 individuals). The experiment took place over 90 days: captive families were placed in the identical tanks to the wild families, but one month later, as wild families were generated one month earlier due to idiosyncrasies between wild and captive spawning times. Captive and wild trout experienced the same tank conditions throughout the experiment (water temperature ∼6.5°C; pH = 7.0, dissolved oxygen = 8.5 mgl^-1^, and daylight between 9:00 and 17:00, cumulative degree days for growth = 800). Wild and captive trout were kept in tubes for 45 days; as fish grew, they were moved to larger tubes in two new (and same sized) tanks for the last 45 days so as not to affect growth. Family replicates within each tank were randomly allocated one of two diet treatments for rearing: drifting prey (*Artemia salina*), hereafter ‘natural diet’ and pellet feed, hereafter referred to as ‘captive/pellet diet. Feeding rates were standardized for first 45 days, 2ml of artemia and 10mg of pellet for each tube. In the last 45 days, the feeding rate was increased to double with 4 ml of artemia. Artemia concentration was standardized by growing 10,00000/L and suspending live artemia from a batch into 2L of water prior to feeding and 20mg of pellet for each tube. As one of our main objectives of this current study was to see the growth and mortality due to exogenous feeding, the feeding frequency was 4 times per day (2 hour intervals) for the first 45 days and then increased to 5 times per day (1.5 hour intervals) for the last 45 days. To ensure complete nutritional needs were met 20mg of pellet feed was given once a day to fish in the artemia treatment for the last 45 days.

### Size, growth and mortality

At the beginning and termination of the experiment, a standardized photo of each individual (wild and captive) was taken using a mounted overhead camera. Photos were then imported into the program ImageJ (Rasband, 2014) and fork length (mm) was measured against a known size standard.

Tubes were closely monitored daily, with dead individuals counted and removed when needed. To maintain the same tube densities (n = 4), a non-experimental individual from the same population was added to a tube when an experimental fish mortality occurred; each non-experimental individual was clipped on the caudal fin to demarcate it from experimental fish. At the termination of the experiment, before measuring the final length, all non-experimental individuals were removed from the tube.

## Statistical Analyses

### Captive and wild size/growth comparisons in relation to diet

General linear mixed effects models (GLMMs) were used to evaluate the effect of population, origin (F1 captive or wild), diet, and their interactions on growth rate with respect to length. Absolute Growth Rate (AGR) was quantified for all fish, which assumes a relatively constant linear rate of growth over the experimental period. Initial and final length were included as dependent variables in a model. Population (FW, CC, WN), origin (captive vs. wild), and diet (artemia vs. pellet) were included as categorical fixed effects. Time was included as a fixed continuous effect; the slope of the time variable represented overall AGR, with the interaction terms between time and the other categorical fixed covariates representing their effect on AGR. All interactions between fixed effects were included. Tank and a tank-by-time interaction were also included as fixed effects to account for the randomized experimental design. A random family intercept and uncorrelated family-by-time term were fitted to the model to account for non-independence among the data due to relatedness of individuals. A random replicate intercept and uncorrelated replicate-by-time term was also included to account for repeated measurements within each replicate. Visual examination of growth trends detected heteroscedastic variation in growth over time among population, origin, and diet: separate variances for each level of time were therefore modelled for each factor-combination level, and likelihood ratio tests (LRT) were used to evaluate the significance of each variance term.

Analysis was conducted using the *nlme* statistical package (Pinheiro et al., 2012) in R 3.5.1 (R Core Team, 2019). Models were estimated using Maximum Likelihood (ML) during model selection; final parameter estimates were obtained under restricted Maximum Likelihood (REML). Non-significant parameters (*p >*0.05) were backwards stepwise removed using LRTs, eliminating higher order interaction terms first. Relevant lower order terms were automatically retained if a higher order term was found to be significant. Tank and a tank-by-date interaction were included in all models regardless of significance to account for the experimental design. Pairwise comparisons of means among significant groupings were conducted using *t*-tests, with degrees of freedom calculations based on the containment method implemented in the r package *emmeans* (v1.3.4, Lenth, 2019), and *p*-values Bonferroni corrected to account for type 1 error rates.

### Captive vs. wild mortality relative to diet

Mortality was analyzed using a generalized linear mixed-effects model with a binomial distribution (logit-link function). The number of surviving and deceased individuals per replicate was modelled as the dependent variable; population, origin, and diet were included as categorical fixed terms. All possible interactions between these terms were tested. The mean initial length of each replicate (centered and scaled) was also included as a fixed continuous covariate to account for potential differences in mortality due to size. Tank was also included as a fixed effect to account for the randomized experimental design. Family was included as a random effect term. Data analysis was conducted using the *lme4* statistical package (Bates et al., 2014) in R 3.5.1 using Laplace approximation to the likelihood. Backwards stepwise model selection and pairwise comparisons were conducted as for the length data.

## Results

### Captive vs. wild size relative to diet

Significant heteroscedastic variation in length over time was detected for all factor-level combinations (Table S2); all variance terms were therefore retained in the final length model. Two three-way interactions between fixed-effects were significant: time-by-population-by-treatment, and source-by-population-by-treatment (Table S3). All relevant lower-order terms were therefore retained. The time-by-source term was highly significant and also retained in the final model.

There were no significant initial length differences between diet treatments within any population-origin factor combination at time 0 due to the randomized assignment of fish to experimental replicates (Table S4). However, all captive-origin fish were substantially larger at time 0 relative to wild-origin fish (Table S5, Figure 1a).

**Fig. 1:**
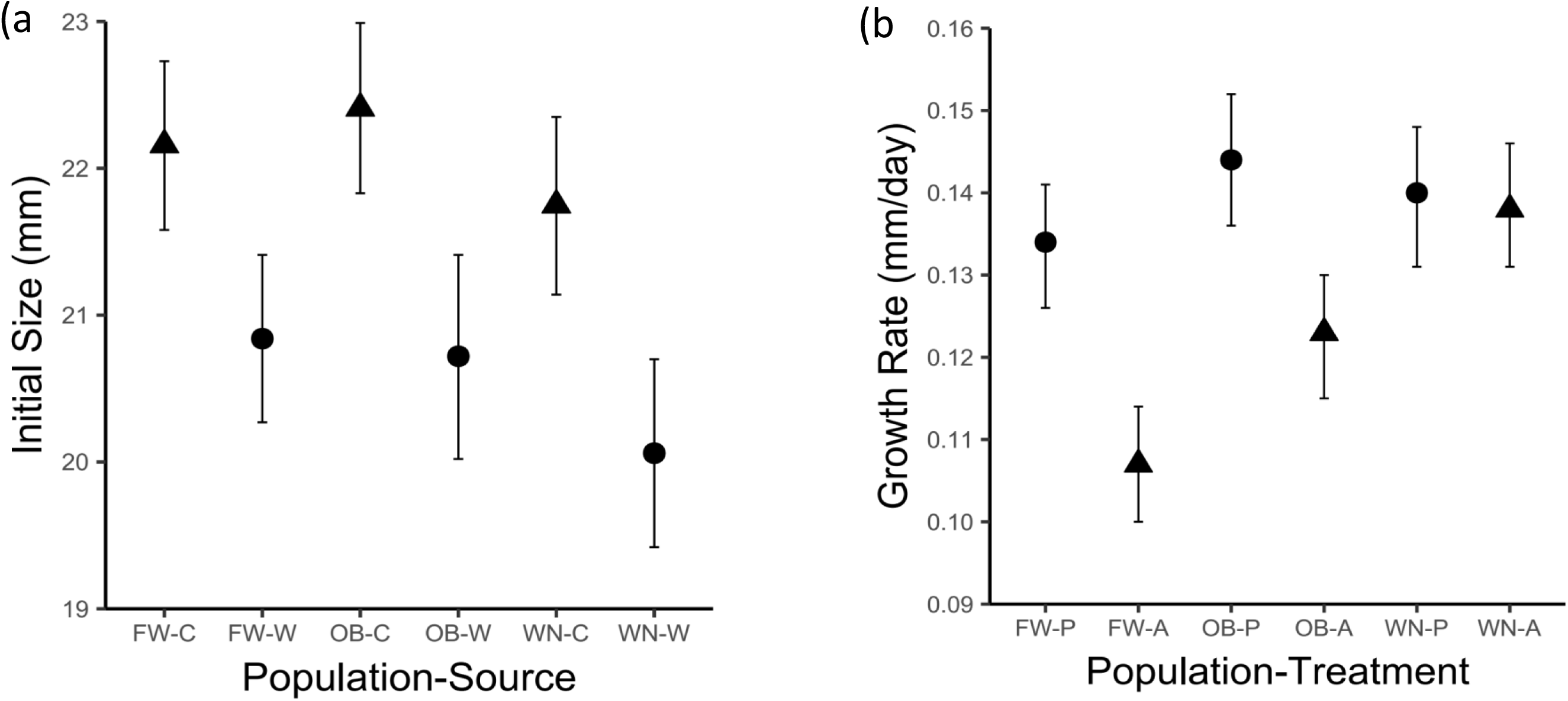
(a) Initial size differences among different populations of captive and wild origin fish; (b) Growth differences among different populations (FW, OB, WN) across treatments (Natural Prey (Artemia = A) and Pellet = P)

Regardless of population of origin or diet treatment, fish from captive-origin parents grew more rapidly than fish from wild-origin parents (*t_1330_* = 3.81; *p* = <0.001, Table S6, Figure 1b). Growth response to diet treatment was consistent across captive and wild fish but differed based on population of origin. Fish from FW and OB grew significantly slower on a natural diet relative to pellets, whereas fish from WN exhibited similar growth regardless of diet treatment (Table 1, Figure 2a). All populations exhibited similar growth on a pellet diet but differed in their relative responses to the natural diet treatment; FW grew significantly slower on the natural diet than OB and WN, and OB grew marginally slower on the natural diet relative to WN (Table 2). A significant treatment-by-source-by-time was not detected (Table S3), indicating both captive and wild fish exhibited similar growth reaction norms across the treatments (Fig. 3).

**Fig. 2:**
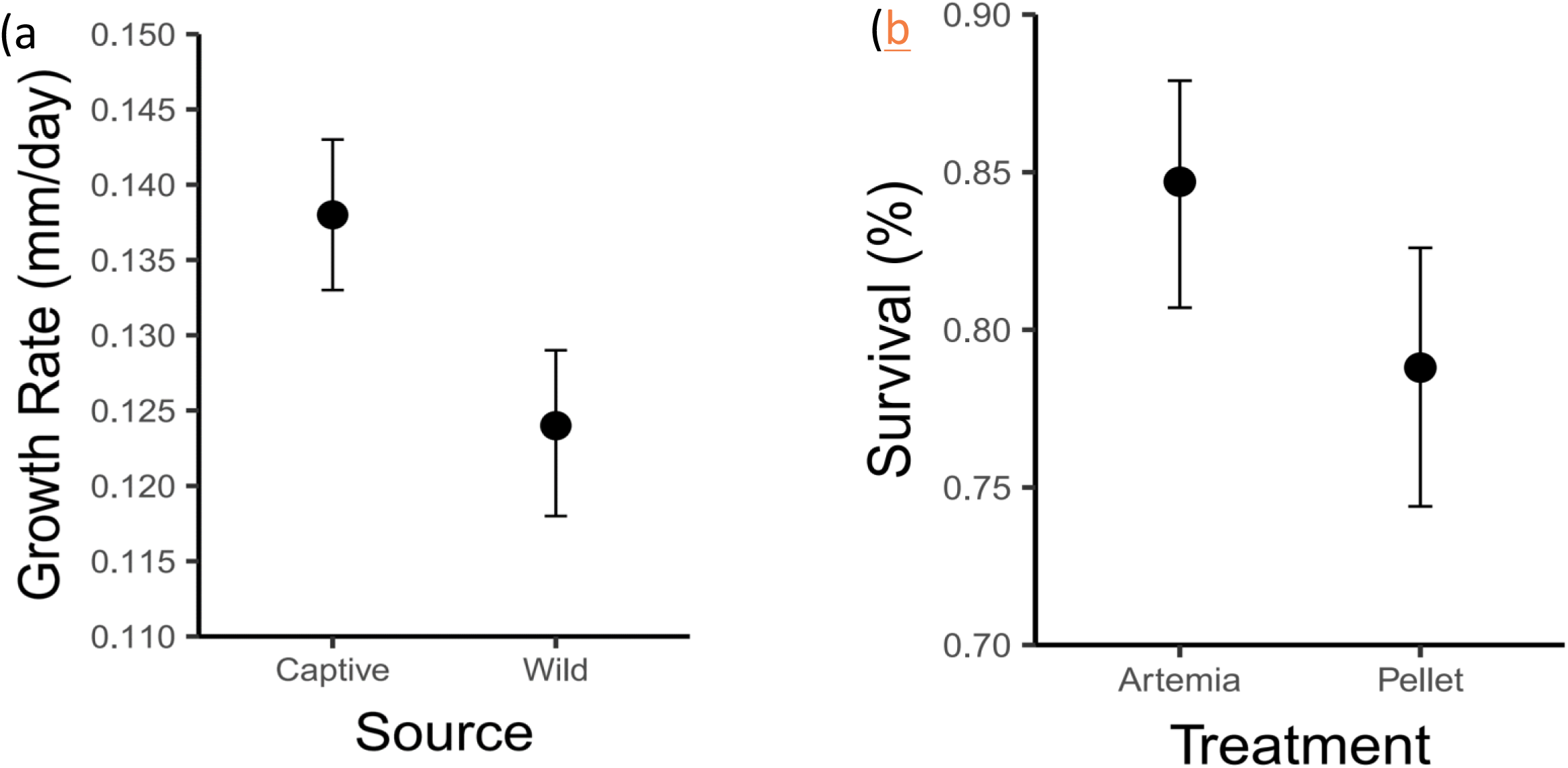
(a) Growth rate differences between captive and wild fish; (b) Survival differences across diet treatments (Natural Prey (Artemia) vs. captive (Pellet))

**Fig. 3:**
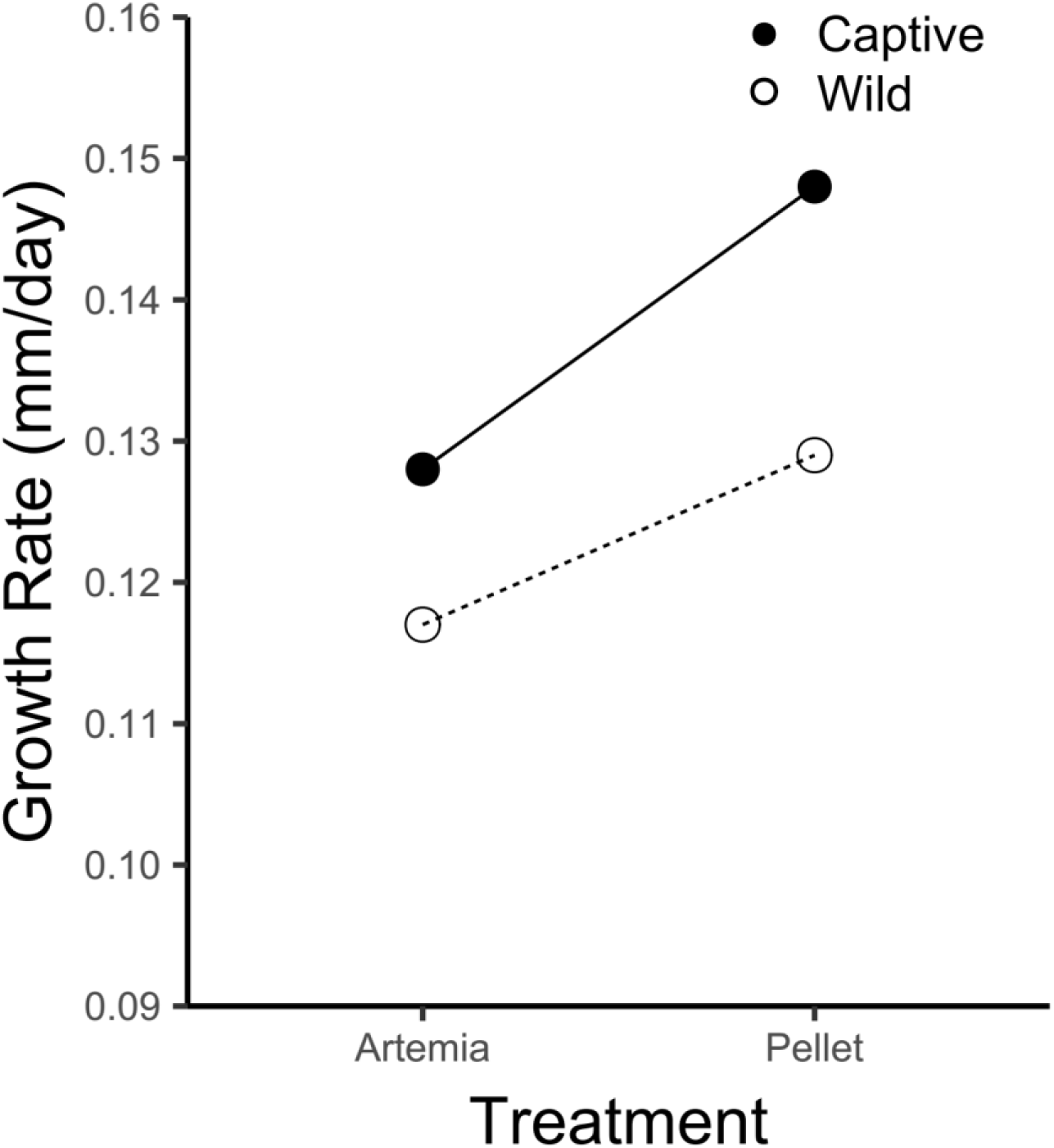
Growth reaction norms for captive and wild origin fish across diet treatments.

**Table 1:** Growth rate differences between populations, within treatments. P-values have been Bonferroni corrected.

| Population contrast | Treatment (diet) | Growth-rate Difference (mm/day) | SE | Df | p-value |
| --- | --- | --- | --- | --- | --- |
| FW - OB | Natural Prey | -0.0157 | 0.0051 | 1330 | 0.019 |
| FW - WN | Natural Prey | -0.0314 | 0.0050 | 1330 | <0.001 |
| OB - WN | Natural Prey | -0.0157 | 0.0053 | 1330 | <0.001 |
| FW - OB | Pellet | -0.0102 | 0.0056 | 1330 | 0.617 |
| FW - WN | Pellet | -0.0060 | 0.0056 | 1330 | 1.000 |
| OB - WN | Pellet | 0.0042 | 0.0058 | 1330 | 1.000 |

**Table 2:**
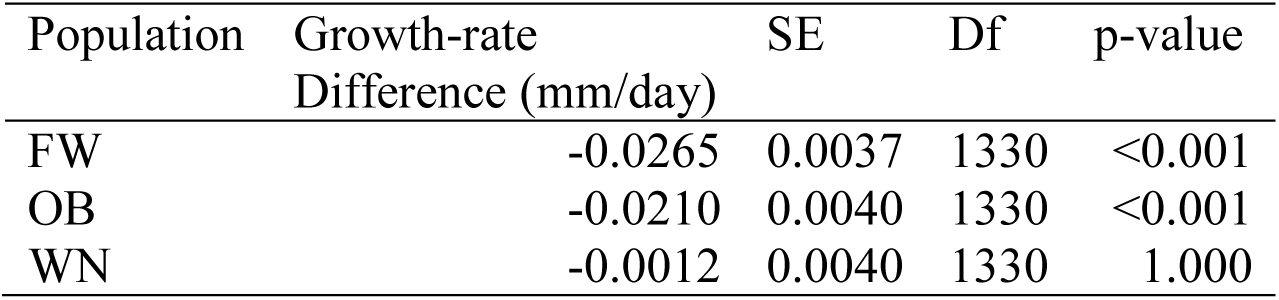
Growth rate differences within population, between treatments (Natural Prey - Pellet). P-values have been Bonferroni corrected.

| Population | Growth-rate<br>Difference (mm/day) | SE | Df | p-value |
| --- | --- | --- | --- | --- |
| FW | -0.0265 | 0.0037 | 1330 | <0.001 |
| OB | -0.0210 | 0.0040 | 1330 | <0.001 |
| WN | -0.0012 | 0.0040 | 1330 | 1.000 |

### Captive vs. wild mortality relative to diet

The only significant term retained after model selection was diet treatment (Table S7). There was no significant difference in mortality between captive and wild fish or amongst the different populations. However, despite exhibiting improved growth in the pellet diet treatment, fish exhibited significantly improved survival in the natural diet treatment (*t_106_* = 2.23, *p* = 0.026, Figure 2b).

## Discussion

Our results support the hypothesis that single generation exposure to a captive diet may act as a primer of domestication selection by inducing genetically-based changes to individual survival and growth in wild animal populations. Namely, under common, captive environmental conditions, we found that (i) early juvenile F_1_ captive brook trout outgrew their wild counterparts regardless of being fed on a captive (pellet) or natural (artemia) diet; (ii) both captive and wild trout had better survival on the natural diet; (iii) populations had an idiosyncratic response to diet treatment; (iv) captive and wild trout exhibited the same response to diet treatments, i.e. there was no difference in plasticity associated with captivity, but (iv) populations differed in their plastic responses to diet.

That captive fish grew more rapidly than wild fish, irrespective of the population of origin or diet treatment, is consistent with observations in other captive-reared salmonid populations (e.g., Campbell et al., 2006; Larsen et al., 2006; Araki et al., 2007; Fraser, 2008; Berejikian et al., 2012). This implies that the pellet diet itself is a selective agent at this early life stage, given that fitness-related traits (e.g., growth) are among the first traits to be affected by intentional or inadvertent domestication selection. For example, high densities in captive environments, combined with feeding regimes that promote competitive behaviours such as dominance and boldness (e.g., repetitive, predictable access to highly nutritious pellet diet) may inadvertently select for growth differences between captive and wild fish (Einum & Fleming, 1997; McGinnity et al., 2003). Depending on the wild environments from which fish originate, behaviours may be fully engrained and inflexible (Huntingford, 2004; Canario et al., 2013; Janisse et al., 2019), resulting in poor adaptation to captivity. Put simply, other traits (e.g., behavioural traits) probably affect growth response in captivity and so it is hard to pin down exactly that only the novel captive diet leads to more rapid growth in captive fish.

We also found that growth responses to captive and natural diets differed among wild populations. While study populations exhibited no difference in their growth on a pellet diet, FW and OB grew faster on a pellet diet than a natural diet, whereas WN grew equally as well on both diets. FW also grew significantly slower on a natural diet than OB and WN; we cannot simply rule out the possibility that FW fish were unable to extract nutritional or caloric value from the natural diet.

Survival was quite high (80-85%) in our study and similar between captive and wild fish but was significantly higher in the natural diet treatment. Based on the observations from a related study where only a pellet diet was adopted (Fraser et al., 2019), we speculate that much of the mortality during rearing took place in the single generation. For example, in that study, lifetime success in captivity among nine Cape Race trout populations was on average 0.15 with a range of 0-0.35 and much of the mortality took place in the first six months of life (lifetime success combined the probability of surviving to maturation for captive-born fish with the early survival probability of their progeny). This suggests that early in life is when much of the opportunity for domestication selection to act occurs. Despite exhibiting higher growth on a captive diet, fish also exhibited poorer survival on this diet. Wild fish had less opportunity to adapt to a novel captive diet given that this was the first time their lineage was exposed to pellet feed. Yet, F_1_ captive families, whose parents were exposed to pellet diet, did not exhibit improved survival relative to wild fish when exposed to a pellet diet. We did not find evidence to suggest that captive fish have become adapted to the form and nutritional content of commercial pellet diet to the extent it influences the survival of their offspring. It is possible that a single generation of selection is not enough to shape diet-based reaction norms for survival.

While we found that our study populations exhibited idiosyncratic plastic responses to diet treatments, these responses were consistent across populations regardless of source. Within a population, diet reaction norms for growth were similar for both captive-bred and wild fish. Likewise, both captive and wild fish exhibited similar survival reaction norms across diet treatments. Comparative studies of phenotypic plasticity in salmonids have revealed differing results (Fraser et al., 2008; Darwish et al., 2009; Morris et al., 2011). Overall, a single generation of domestication might not be enough to result in genetic changes to reaction norms, given that both captive and wild fish in our study originated from the same genetic/phenotypic starting conditions.

In summary, we find evidence that a single generation exposure to a captive diet generates genetically-based changes to growth but not survival in a wild fish. Captive diets may therefore contribute to and prime maladaptation generated in wild species from captive rearing. However, a single generation might not be enough to shape adaptation in some traits to a commercial diet. Understanding the impacts of a captive diet on phenotypic and genetic differences between captive and wild animals therefore merits further attention in conservation and supplementation of wild species of conservation concern, in addition to the sustainable development of captive rearing and supplementation. Our research further supports the contention that the scope of maladaptive effects to wild fitness from a single generation of captive exposure may vary considerably among intraspecific populations (Fraser et al., 2019) and should be taken into account when implementing species conservation programs.

## Acknowledgements

This research was supported by NSERC Discovery and Accelerator Grants to DJF, NSERC graduate scholarships to MCY, an Erasmus Mundus Scholarship to SSI and by the European Commission to SSI, and was carried out in accordance with the guidelines set by the Concordia Animal Research Ethics Committee.

---

**Table S1:** Number of individuals and number of families per wild and captive populations used in the experiment

| Population | Source | Treatment | Tank | Family | Individuals |
| --- | --- | --- | --- | --- | --- |
| FW | Captive | Artemia | 1 | 10 | 40 (10 x 4) |
| FW | Captive | Artemia | 2 | 10 | 40 (10 x 4) |
| FW | Captive | Pellet | 1 | 10 | 39 (9 x 4; 1 x 3) |
| FW | Captive | Pellet | 2 | 10 | 40 (10 x 4) |
| FW | Wild | Artemia | 1 | 10 | 40 (10 x 4) |
| FW | Wild | Artemia | 2 | 10 | 40 (10 x 4) |
| FW | Wild | Pellet | 1 | 10 | 40 (10 x 4) |
| FW | Wild | Pellet | 2 | 10 | 38 (8 x 4; 2 x 3) |
| OB | Captive | Artemia | 1 | 10 | 40 (10 x 4) |
| OB | Captive | Artemia | 2 | 10 | 40 (10 x 4) |
| OB | Captive | Pellet | 1 | 10 | 40 (10 x 4) |
| OB | Captive | Pellet | 2 | 10 | 40 (10 x 4) |
| OB | Wild | Artemia | 1 | 7 | 28 (7 x 4) |
| OB | Wild | Artemia | 2 | 7 | 28 (7 x 4) |
| OB | Wild | Pellet | 1 | 7 | 28 (7 x 4) |
| OB | Wild | Pellet | 2 | 7 | 28 (7 x 4) |
| WN | Captive | Artemia | 1 | 9 | 35 (8 x 4; 1 x 3) |
| WN | Captive | Artemia | 2 | 9 | 36 (9 x 4) |
| WN | Captive | Pellet | 1 | 9 | 36 (9 x 4) |
| WN | Captive | Pellet | 2 | 9 | 36 (9 x 4) |
| WN | Wild | Artemia | 1 | 8 | 32 (8 x 4) |
| WN | Wild | Artemia | 2 | 8 | 32 (8 x 4) |
| WN | Wild | Pellet | 1 | 8 | 32 (8 x 4) |
| WN | Wild | Pellet | 2 | 8 | 31 (7 x 4; 1 x 3) |
|  |  |  |  |  | Total No: 859 |

**Table S2:**
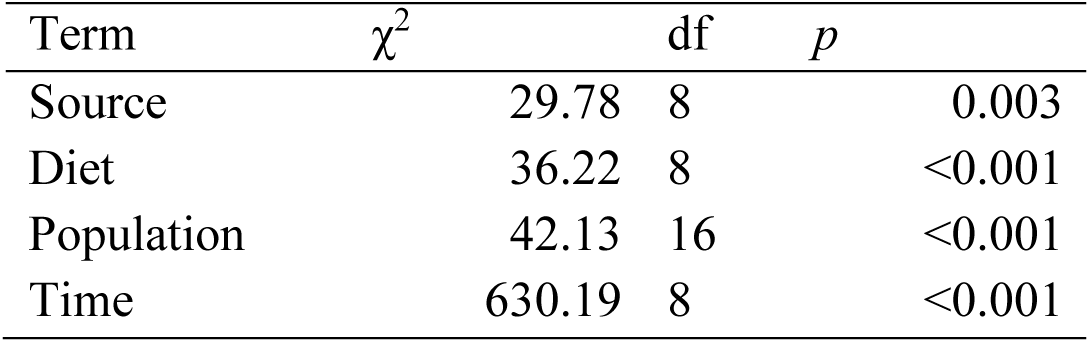
Results of Likelihood ratio tests evaluating the significance of heteroscedastic variance terms in growth model.

**Table S3:** Results of model selection for growth analysis, with statistically significant (retained) terms presented in bold. Model testing was conducted using LRTs.

| Model No. | Description | Versus model No. | Term | $\chi^2$ | df | *p* |
| --- | --- | --- | --- | --- | --- | --- |
| 0 | Tank + Treatment + Pop + Source + Time + Tank:Time + Treatment:Pop + Treatment:Source + Treatment:Time + Pop:Source + Pop:Time + Source:Time + Treatment:Pop:Source + Treatment:Pop:Time + Treatment:Source:Time + Pop:Source:Time + Treatment:Pop:Source:Time | - |  |  |  |  |
| 1 | Tank + Treatment + Pop + Source + Time + Tank:Time + Treatment:Pop + Treatment:Source + Treatment:Time + Pop:Source + Pop:Time + Source:Time + Treatment:Pop:Source + Treatment:Pop:Time + Treatment:Source:Time + Pop:Source:Time | 0 | Treatment:<br>Pop:Source:<br>Time | 2.64 | 2 | 0.267 |
| 2 | Tank + Treatment + Pop + Source + Time + Tank:Time + Treatment:Pop + Treatment:Source + Treatment:Time + Pop:Source + Pop:Time + Source:Time + Treatment:Pop:Source + Treatment:Pop:Time + Treatment:Source:Time | 1 | Pop:Source:<br>Time | 3.34 | 2 | 0.188 |
| †3 | Tank + Treatment + Pop + Source + Time + Tank:Time + Treatment:Pop + Treatment:Source + Treatment:Time + Pop:Source + Pop:Time + Source:Time + Treatment:Pop:Source + Treatment:Pop:Time | 2 | Treatment:<br>Source:Time | 2.72 | 1 | 0.099 |
| 4 | Tank + Treatment + Pop + Source + Time + Tank:Time + Treatment:Pop + Treatment:Source + Treatment:Time + Pop:Source + Pop:Time + Source:Time | 3 | **Treatment:**<br>**Pop:Source** | 9.48 | 2 | **0.009** |
| 5 | Tank + Treatment + Pop + Source + Time + Tank:Time + Treatment:Pop + Treatment:Source + Treatment:Time + Pop:Source + Pop:Time + Source:Time | 3 | **Treatment:**<br>**Pop:Time** | 22.37 | 2 | **<0.001** |
| 6 | Tank + Treatment + Pop + Source + Time + Tank:Time + Treatment:Pop + Treatment:Source + Treatment:Time + Pop:Source + Pop:Time + Treatment:Pop:Source + Treatment:Pop:Time | 3 | **Source:Time** | 12.96 | 1 | **<0.001** |

†Retained model

**Table S4:**
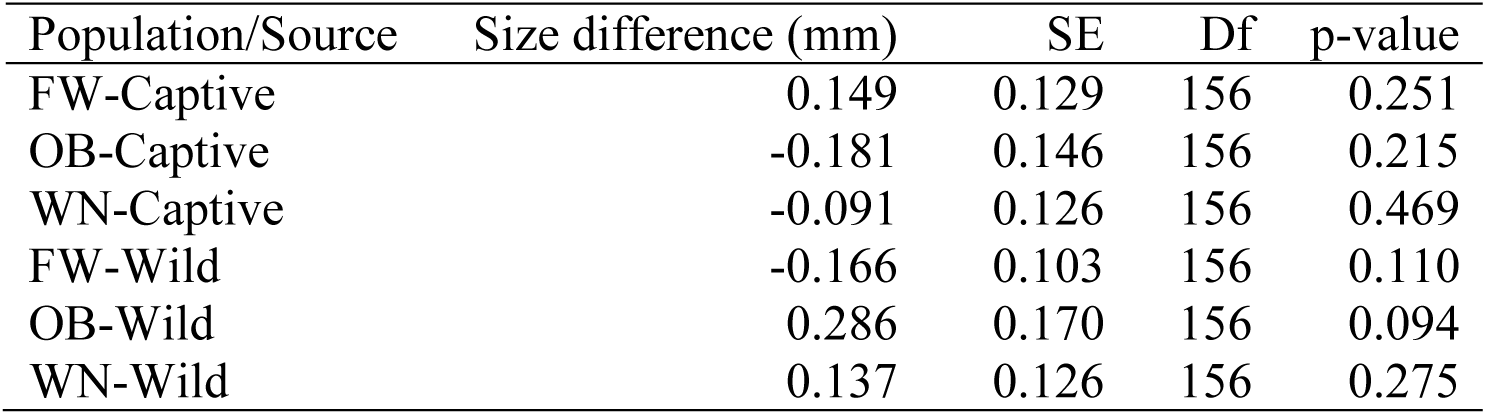
Initial size difference comparisons between natural prey and pellet treatments within a population and source (Natural prey (Artemia) treatments are the reference group). P-values have been Bonferroni corrected.

**Table S5:** Initial size differences (Captive - Wild), by population.

| Population | Size difference (mm) | SE | Df | p-value |
| --- | --- | --- | --- | --- |
| FW | 1.314 | 0.403 | 48 | 0.002 |
| OB | 1.690 | 0.449 | 48 | <0.001 |
| WN | 1.691 | 0.438 | 48 | <0.001 |

**Table S6:** Growth rate differences between sources (captive - wild)

| Growth difference (mm/day) | SE | Df | p-value |
| --- | --- | --- | --- |
| 0.0145 | 0.0038 | 1330 | <0.001 |

**Table S7:**
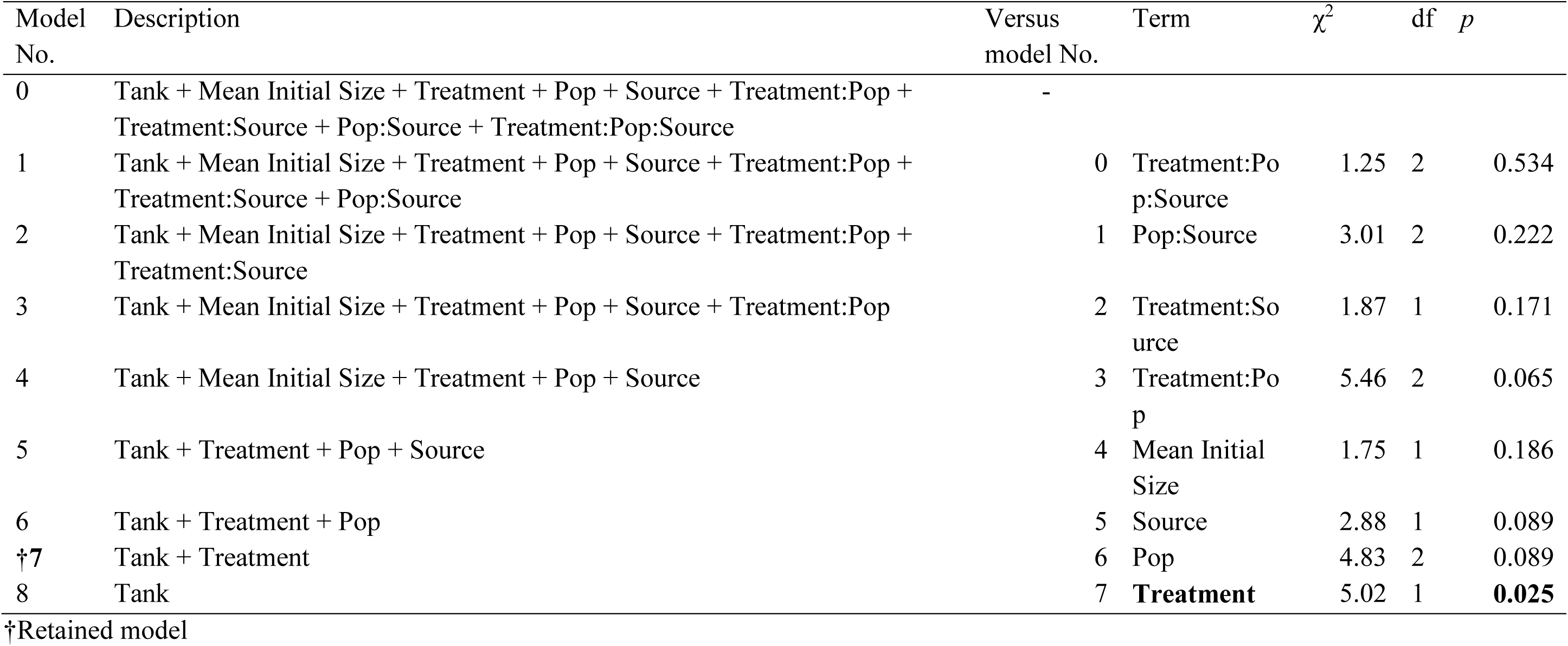
Results of model selection for survival analysis, with statistically significant (retained) terms presented in bold. Model testing was conducted using LRTs.

**Fig. S1:**
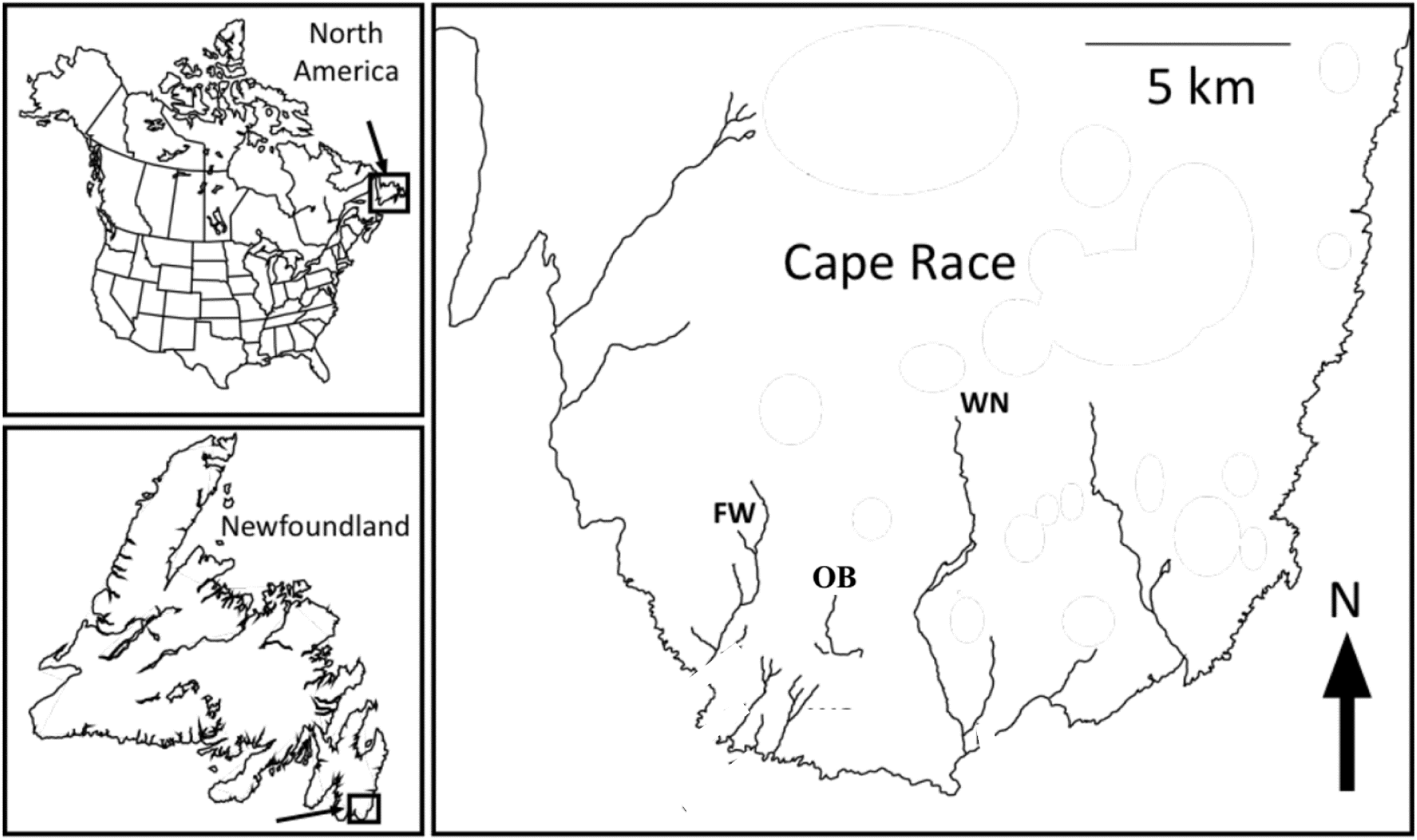
The study site in Cape Race, Newfoundland, Canada (FW=Freshwater; WN=Watern cove; OB=Upper Ouananiche Beck) (Adapted from: Wood, J. L. (2014). (Doctoral dissertation, Concordia University, Montreal, Canada).

## References

1. Allan, N., Knotts, T., Pesapane, R., Ramsey, J., Castle, S., Clifford, D., & Foley, J. (2018). Conservation Implications of Shifting Gut Microbiomes in Captive-Reared Endangered Voles Intended for Reintroduction into the Wild. Microorganisms 6, 94.

2. Araki, H., Cooper, B., & Blouin, M. S. (2007). Genetic effects of captive breeding cause a rapid, cumulative fitness decline in the wild. Science. 318, 100–103.

3. Araki, H., Cooper, B., & Blouin, M. S. (2009). Carry-over effect of captive breeding reduces reproductive fitness of wild-born descendants in the wild. Biol. Lett. 5, 621–624.

4. Baskett, M. L., & Waples, R. S. (2013). Evaluating alternative strategies for minimizing unintended fitness consequences of cultured individuals on wild populations. Conserv. Biol. 27, 83–94.

5. Bates, D., Mächler, M., Bolker, B., & Walker, S. (2014). Fitting linear mixed-effects models using lme4. arXiv Prepr. arXiv1406.5823.

6. Benhaïm, D., Guyomard, R., Chatain, B., Quillet, E., & Bégout, M.-L. (2013). Genetic differences for behaviour in juveniles from two strains of brown trout suggest an effect of domestication history. Appl. Anim. Behav. Sci. 147, 235–242.

7. Berejikian, B. A., Larsen, D. A., Swanson, P., Moore, M. E., Tatara, C. P., Gale, W. L., Pasley, C. R., & Beckman, B. R. (2012). Development of natural growth regimes for hatchery-reared steelhead to reduce residualism, fitness loss, and negative ecological interactions. Environ. Biol. Fishes 94, 29–44.

8. Bernos, T. A., & Fraser, D. J. (2016). Spatiotemporal relationship between adult census size and genetic population size across a wide population size gradient. Mol. Ecol. 25, 4472–4487.

9. Bowlby, H. D., & Gibson, A. J. F. (2011). Reduction in fitness limits the useful duration of supplementary rearing in an endangered salmon population. Ecol. Appl. 21, 3032–3048.

10. Braithwaite, V. A., & Salvanes, A. G. V. (2005). Environmental variability in the early rearing environment generates behaviourally flexible cod: implications for rehabilitating wild populations. Proc. R. Soc. B Biol. Sci. 272, 1107–1113.

11. Brightsmith, D. J. (2012). Nutritional levels of diets fed to captive Amazon parrots: does mixing seed, produce, and pellets provide a healthy diet? J. Avian Med. Surg. 26, 149–161.

12. Campbell, B., Beckman, B. R., Fairgrieve, W. T., Dickey, J. T., & Swanson, P. (2006). Reproductive investment and growth history in female coho salmon. Trans. Am. Fish. Soc. 135, 164–173.

13. Canario, L., Mignon-Grasteau, S., Dupont-Nivet, M., & Phocas, F. (2013). Genetics of behavioural adaptation of livestock to farming conditions. Animal 7, 357–377.

14. Carciofi, A. C., Duarte, J. M. B., Mendes, D., & de Oliveira, L. D. (2006). Food selection and digestibility in yellow-headed conure (Aratinga jandaya) and golden-caped conure (Aratinga auricapilla) in captivity. J. Nutr. 136, 2014S–2016S.

15. Christie, M. R., Marine, M. L., French, R. A., & Blouin, M. S. (2012). Genetic adaptation to captivity can occur in a single generation. Proc. Natl. Acad. Sci. 109, 238–242.

16. Christie, M. R., Marine, M. L., Fox, S. E., French, R. A., & Blouin, M. S. (2016). A single generation of domestication heritably alters the expression of hundreds of genes. Nat. Commun. 7, 10676.

17. Christie, M. R., McNickle, G. G., French, R. A., & Blouin, M. S. (2018). Life history variation is maintained by fitness trade-offs and negative frequency-dependent selection. Proc. Natl. Acad. Sci. 115, 4441–4446.

18. Clarke, C. N., Fraser, D. J., & Purchase, C. F. (2016). Lifelong and carry-over effects of early captive exposure in a recovery program for Atlantic salmon (Salmo salar). Anim. Conserv. 19, 350–359.

19. Darwish, T. L., & Hutchings, J. A. (2009). Genetic variability in reaction norms between farmed and wild backcrosses of Atlantic salmon (Salmo salar). Can. J. Fish. Aquat. Sci. 66, 83–90.

20. De Mestral, L. G., & Herbinger, C. M. (2013). Reduction in antipredator response detected between first and second generations of endangered juvenile Atlantic salmon Salmo salar in a captive breeding and rearing programme. J. Fish Biol. 83, 1268–1286.

21. Dender, M. G. E., Capelle, P. M., Love, O. P., Heath, D. D., Heath, J. W., & Semeniuk, C. A. D. (2018). Phenotypic integration of behavioural and physiological traits is related to variation in growth among stocks of Chinook salmon. Can. J. Fish. Aquat. Sci. 75, 2271–2279.

22. DeWitt, T. J., & Scheiner, S. M. (2004). Phenotypic plasticity: functional and conceptual approaches. Oxford University Press.

23. Edelaar, P., & Bolnick, D. I. (2019). Appreciating the multiple processes increasing individual or population fitness. Trends Ecol. Evol. 34, 435–446.

24. Erlenbach, J. A., Rode, K. D., Raubenheimer, D., & Robbins, C. T. (2014). Macronutrient optimization and energy maximization determine diets of brown bears. J. Mammal. 95, 160–168.

25. Einum, S., & Fleming, I. A. (1997). Genetic divergence and interactions in the wild among native, farmed and hybrid Atlantic salmon. J. Fish Biol. 50, 634–651.

26. Evans, M. L., Wilke, N. F., O’Reilly, P. T., & Fleming, I. A. (2014). Transgenerational effects of parental rearing environment influence the survivorship of captive-born offspring in the wild. Conserv. Lett. 7, 371–379.

27. Fleming, I. A., & Petersson, E. (2001). The ability of released, hatchery salmonids to breed and contribute to the natural productivity of wild populations. Nord. J. Freshw. Res. 71–98.

28. Ford, M. J. (2002). Selection in captivity during supportive breeding may reduce fitness in the wild. Conserv. Biol. 16, 815–825.

29. Frankham, R. (2008). Genetic adaptation to captivity in species conservation programs. Mol. Ecol. 17, 325–333.

30. Fraser, D. J. (2008). How well can captive breeding programs conserve biodiversity? A review of salmonids. Evol. Appl. 1, 535–586.

31. Fraser, D. J., Cook, A. M., Eddington, J. D., Bentzen, P., & Hutchings, J. A. (2008). Mixed evidence for reduced local adaptation in wild salmon resulting from interbreeding with escaped farmed salmon: complexities in hybrid fitness. Evol. Appl. 1, 501–512.

32. Fraser, D. J., Debes, P. V, Bernatchez, L., & Hutchings, J. A. (2014). Population size, habitat fragmentation, and the nature of adaptive variation in a stream fish. Proc. R. Soc. B Biol. Sci. 281, 20140370.

33. Fraser, D. J., Walker, L., Yates, M. C., Marin, K., Wood, J. L. A., Bernos, T. A., & Zastavniouk, C. (2019). Population correlates of rapid captive-induced maladaptation in a wild fish. Evol. Appl. 12, 1305–1317.

34. Gozlan, R. E., Britton, J. R., Cowx, I., & Copp, G. H. (2010). Current knowledge on non-native freshwater fish introductions. J. Fish Biol. 76, 751–786.

35. Hagen, I. J., Jensen, A. J., Bolstad, G. H., Diserud, O. H., Hindar, K., Lo, H., & Karlsson, S. (2019). Supplementary stocking selects for domesticated genotypes. Nat. Commun. 10, 199.

36. Hewson-Hughes, A. K., Colyer, A., Simpson, S. J., & Raubenheimer, D. (2016). Balancing macronutrient intake in a mammalian carnivore: disentangling the influences of flavour and nutrition. R. Soc. open Sci. 3, 160081.

37. Huntingford, F. A. (2004). Implications of domestication and rearing conditions for the behaviour of cultivated fishes. J. Fish Biol. 65, 122–142.

38. Hutchings, J. A. (1993). Reaction norms for reproductive traits in brook trout and their influence on life history evolution affected by size-selective harvesting. In Exploit. Evol. Resour. pp. 107–125. Springer.

39. Janisse, K., Capelle, P. M., Heath, J. W., Dender, M. G. E., Heath, D. D., & Semeniuk, C. A. D. (2019). Life in captivity: Varied behavioural responses to novel setting and food types in first-generation hybrids of farmed and wild juvenile Chinook salmon (Oncorhynchus tshawytscha). Can. J. Fish. Aquat. Sci.76, 1962–1970

40. Jonsson, B., & Jonsson, N. (2011). Habitats as Template for Life Histories. In Ecol. Atl. Salmon Brown Trout. pp. 1–21. Springer.

41. Kyriazakis, I., Emmans, G. C., & Whittemore, C. T. (1991). The ability of pigs to control their protein intake when fed in three different ways. Physiol. Behav. 50, 1197–1203.

42. Lall, S. P., & Dumas, A. (2015). Nutritional requirements of cultured fish: Formulating nutritionally adequate feeds. In Feed Feed. Pract. Aquac. pp. 53–109. Elsevier.

43. Larsen, D. A., Beckman, B. R., Strom, C. R., Parkins, P. J., Cooper, K. A., Fast, D. E., & Dickhoff, W. W. (2006). Growth modulation alters the incidence of early male maturation and physiological development of hatchery-reared spring Chinook salmon: a comparison with wild fish. Trans. Am. Fish. Soc. 135, 1017–1032.

44. Le Boucher, R., Dupont-Nivet, M., Vandeputte, M., Kerneis, T., Goardon, L., Labbe, L., Chatain, B., Bothaire, M. J., Larroquet, L., & Medale, F. (2012). Selection for adaptation to dietary shifts: towards sustainable breeding of carnivorous fish. PLoS One 7, e44898.

45. Le Cam, S., Perrier, C., Besnard, A.-L., Bernatchez, L., & Evanno, G. (2015). Genetic and phenotypic changes in an Atlantic salmon population supplemented with non-local individuals: a longitudinal study over 21 years. Proc. R. Soc. B Biol. Sci. 282, 20142765.

46. Lenth, R. (2019). Emmeans: Estimated marginal means, aka least-squares means. Retrieved from https://CRAN.R-project.org/package=emmeans

47. Li, M. H., & Robinson, E. H. (2015). Complete feeds—intensive systems. In Feed Feed. Pract. Aquac. pp. 111–126. Elsevier.

48. Mayntz, D., Raubenheimer, D., Salomon, M., Toft, S., & Simpson, S. J. (2005). Nutrient-specific foraging in invertebrate predators. Science. 307, 111–113.

49. Mayntz, D., Nielsen, V. H., Sørensen, A., Toft, S., Raubenheimer, D., Hejlesen, C., & Simpson, S. J. (2009). Balancing of protein and lipid intake by a mammalian carnivore, the mink, Mustela vison. Anim. Behav. 77, 349–355.

50. McGinnity, P., Prodöhl, P., Ferguson, A., Hynes, R., Maoiléidigh, N. ó, Baker, N., Cotter, D., O’Hea, B., Cooke, D., & Rogan, G. (2003). Fitness reduction and potential extinction of wild populations of Atlantic salmon, Salmo salar, as a result of interactions with escaped farm salmon. Proc. R. Soc. London. Ser. B Biol. Sci. 270, 2443–2450.

51. Morris, M. R. J., Fraser, D. J., Eddington, J., & Hutchings, J. A. (2011). Hybridization effects on phenotypic plasticity: experimental compensatory growth in farmed-wild Atlantic salmon. Evol. Appl. 4, 444–458.

52. Naish, K. A., Taylor III, J. E., Levin, P. S., Quinn, T. P., Winton, J. R., Huppert, D., & Hilborn, R. (2007). An evaluation of the effects of conservation and fishery enhancement hatcheries on wild populations of salmon. Adv. Mar. Biol. 53, 61–194.

53. Näslund, J., & Johnsson, J. I. (2016). Environmental enrichment for fish in captive environments: effects of physical structures and substrates. Fish Fish. 17, 1–30.

54. Pinheiro, J., Bates, D., DebRoy, S., Sarkar, D., & Team, R. C. (2012). nlme: Linear and nonlinear mixed effects models. R Packag. version 3.

55. R Core Team (2019). R: A language and environment for statistical computing. R Foundation for Statistical Computing, Vienna, Austria. Retrieved from http://www.R-project.org/.

56. Rasband, W., (2014). National Institutes of Health, USA. Retrieved from http://imagej.nih.gov/ij.

57. Reed, T. E., Schindler, D. E., & Waples, R. S. (2011). Interacting effects of phenotypic plasticity and evolution on population persistence in a changing climate. Conserv. Biol. 25, 56–63.

58. Reisenbichler, R. R., & Rubin, S. P. (1999). Genetic changes from artificial propagation of Pacific salmon affect the productivity and viability of supplemented populations. ICES J. Mar. Sci. 56, 459–466.

59. Reisenbichler, R. R., Rubin, S., Wetzel, L., & Phelps, S. (2004). Natural selection after release from a hatchery leads to domestication in steelhead. Oncorhynchus mykiss 371–384.

60. Solberg, M. F., Zhang, Z., & Glover, K. A. (2015). Are farmed salmon more prone to risk than wild salmon? Susceptibility of juvenile farm, hybrid and wild Atlantic salmon Salmo salar L. to an artificial predator. Appl. Anim. Behav. Sci. 162, 67–80.

61. Waters, C. D., Hard, J. J., Brieuc, M. S. O., Fast, D. E., Warheit, K. I., Knudsen, C. M., Bosch, W. J., & Naish, K. A. (2018). Genomewide association analyses of fitness traits in captive-reared Chinook salmon: Applications in evaluating conservation strategies. Evol. Appl. 11, 853–868.

62. Wood, J. L. A., Belmar-Lucero, S., Hutchings, J. A., & Fraser, D. J. (2014). Relationship of habitat variability to population size in a stream fish. Ecol. Appl. 24, 1085–1100.

63. Wood, J. L. A., & Fraser, D. J. (2015). Similar plastic responses to elevated temperature among different-sized brook trout populations. Ecology 96, 1010–1019.

64. Wood, J. L. A., Tezel, D., Joyal, D., & Fraser, D. J. (2015). Population size is weakly related to quantitative genetic variation and trait differentiation in a stream fish. Evolution (N. Y). 69, 2303–2318.

65. Woodworth, L. M., Montgomery, M. E., Briscoe, D. A., & Frankham, R. (2002). Rapid genetic deterioration in captive populations: causes and conservation implications. Conserv. Genet. 3, 277–288.

66. Wringe, B. F., Purchase, C. F., & Fleming, I. A. (2016). In search of a “cultured fish phenotype”: a systematic review, meta-analysis and vote-counting analysis. Rev. fish Biol. Fish. 26, 351– 373.

